# Nuclear Receptor Liver Receptor Homolog-1 drives Intestinal Stem Cell regeneration and UPR/ER stress response after gut injury

**DOI:** 10.64898/2026.05.15.725547

**Authors:** Hsi-Ju Chen, Kristina N Braverman, Bohan Yu, James R Bayrer

## Abstract

Stem cell renewal and crypt survival are tightly controlled processes critical for gut repair. Defining key regulators of intestinal healing is critical for the development of new epithelial-targeted therapies. We previously showed that the nuclear receptor LRH-1 (NR5A2) maintains intestinal epithelial health and protects against inflammatory damage. Here, using lineage tracing and selective LRH-1 knockout in the Atoh1^+^ secretory lineage we show LRH-1 is vital for intestinal stem cell (ISC) regeneration in complementary in vivo and ex vivo injury-recovery models. Transcriptomic profiling and pathway analysis reveal downregulation of ER stress and unfolded protein response (UPR) programs. Using a new in vivo model to ascertain how LRH-1 directly impacts intestinal cell responses, we identify key ER stress response genes Ire1α and Xbp1 as potential LRH-1 targets. Together our results uncover a novel mechanism whereby LRH-1 sustains the IRE1α-XBP1 arm of the UPR to support injury-induced dedifferentiation and ISC regeneration. Our findings highlight LRH-1 as a promising therapeutic target for restoring epithelial integrity in inflammatory intestinal disorders.

## Introduction

Liver Receptor Homolog-1 (LRH-1, NR5A2) is an atypical nuclear receptor important for intestinal epithelial renewal^1-3^. Overexpression or pharmacological activation of LRH-1 ameliorates intestinal inflammation in animal models of colitis and TNFα-induced cellular injury in organoids, whereas loss of LRH-1 leads to cell death and an intensified inflammatory response^4-6^. Potential roles for LRH-1 in extramedullary corticosteroid synthesis and antiapoptotic pathways have been proposed to account for these observations^7^. Intriguingly, LRH-1 functions as a pioneering and pluripotency factor in other cellular contexts^8-10^, raising the possibility that this receptor may also facilitate epithelial dedifferentiation and ISC repopulation during injury-induced repair.

The intestinal epithelium comprises an immense exposed surface area performing essential digestive, metabolic, and immunological functions that rely on barrier integrity^11^. The intestinal epithelial cells that line the lumen of the gut undergo continuous renewal in a tightly controlled process powered by *leucine-rich repeat receptor* (*Lgr5*) expressing intestinal stem cells (ISCs) and their progenitors^12,13^. Progenitor cells rapidly expand and commit to either the secretory lineage marked by Atonal-1 (Atoh1) or the absorptive lineage marked by Notch-1^14^.

Following intestinal injury and loss of Lgr5^+^ ISCs, Lgr5^+^ ISC-independent repair mechanisms are activated to restore tissue integrity^15-19^. Multiple studies have demonstrated that intestinal epithelial cells (IECs) committed to the secretory lineages, marked by Atoh1 or other secretory lineage genes, can dedifferentiate, repopulate lost ISCs, and renew the epithelial lining^15,19^. Similarly, cells within the absorptive enterocyte lineage share this capacity for dedifferentiation and can regenerate ISCs after injury^18^. The molecular mechanisms regulating dedifferentiation and epithelial repair, however, are incompletely understood.

In this study, we investigate a potential role for LRH-1 in gut epithelial repair and ISC regeneration using a combination of in vivo and ex vivo damage and recovery models. Using lineage tracing, transcriptomics, and interrogation of LRH-1 chromatin binding we uncover an unexpected mechanism whereby LRH-1 leverages the ER stress response pathway to promote cellular resilience to injury that we propose protects epithelial cells undergoing dedifferentiation. Our findings highlight LRH-1 as a promising therapeutic target to treat intestinal disorders like inflammatory bowel disease (IBD).

## Results

### *Lrh1* is robustly expressed in small intestinal crypts

Previously, cell-specific characterization of *Lrh1* expression in the small intestine has been technically limited owing to inadequate LRH-1-directed antibodies. Single-molecular RNA fluorescence in situ hybridization (SmRNA FISH, RNAscope) on mouse small intestine tissue demonstrates robust *Lrh1* expression in the transient amplifying zone comprised by both absorptive and secretory progenitors (Fig. 1A-C). *Lrh1* expression further overlaps with ISC marker *Olfm4* and the secretory progenitor marker *Atoh1* (Fig. 1B,C). Outside of the intestinal crypts, *Lrh1* expression continues to be strongly detected throughout the mid-villus region with scattered expression persisting into the villus tips, suggesting broader expression than previously described^1,4^ (Fig. 1A). Based on the LRH-1 intestinal epithelial expression pattern and importance in colitis^4-6^, we decided to investigate two distinct potential roles for this receptor in 1) lineage maintenance and 2) stem cell regeneration following intestinal injury.

**Figure 1.**
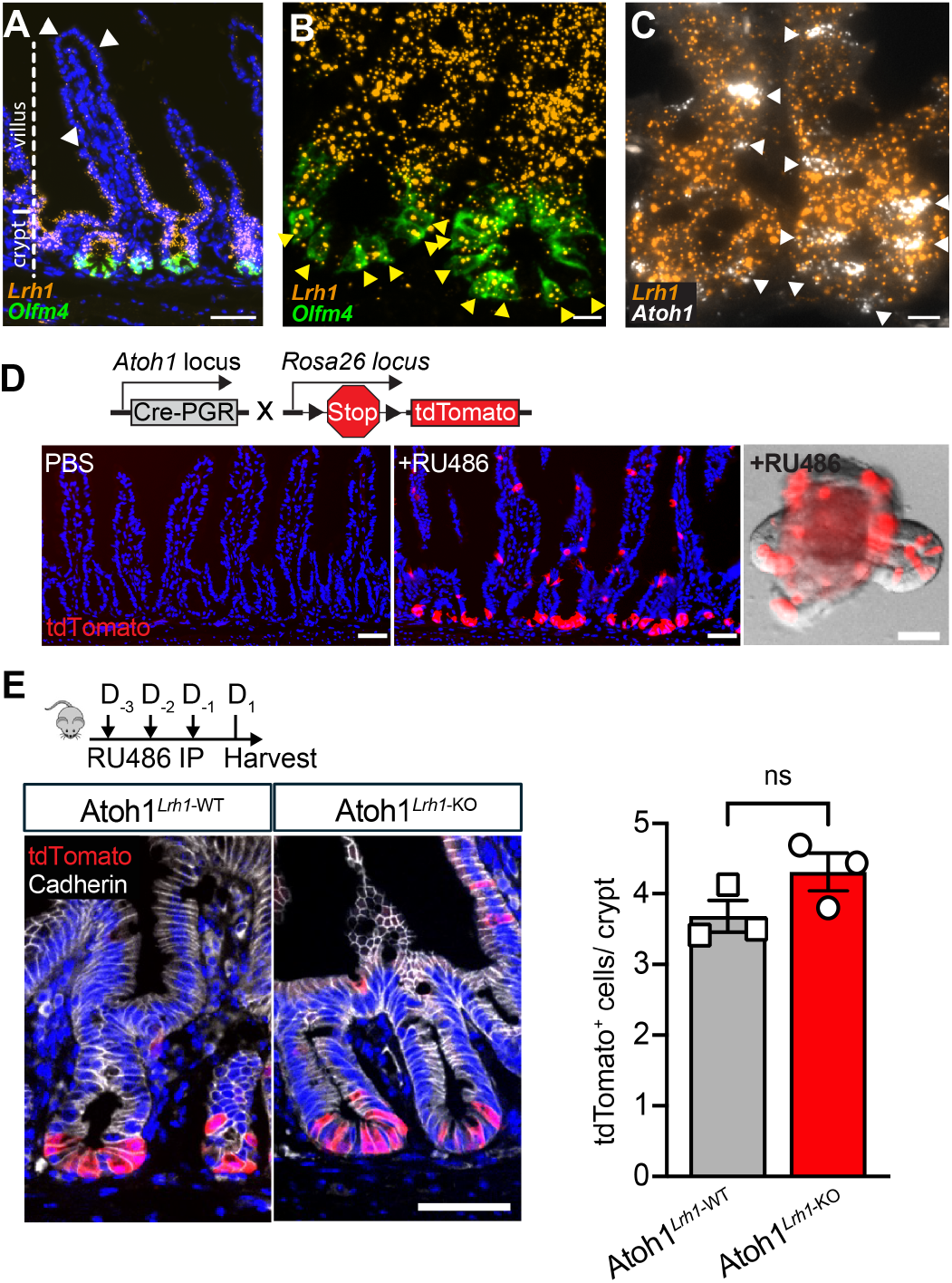
Atoh1^CrePRG^ *Lrh1* knockout does not impair intestinal crypt homeostasis. **(A)** Spatial expression of *Lrh1* and *Olfm4* as detected by RNAscope in mouse small intestine. *Olfm4* is detected only in the crypt. *Lrh1* expression is robust in the crypt but sparse expression in villi (white arrowheads). Scale bar is 50μm. **(B)** Co-localization of *Lrh1* and *Olfm4* in the crypt base ISCs (yellow arrowhead). Scale bar is 10μm. **(C)** Co-localization of *Lrh1* and *Atoh1* (white arrowhead). Scale bar is10μm. **(D)** Schematic diagram illustrating the *Atoh1*^CrePGR^-mediated lineage labeling and representative images from mouse *in vivo* intestinal epithelium and *ex vivo* intestinal organoid. Scale bar is 50μm. **(E)** Representative images and quantification show *Atoh1*^CrePGR^-mediated acute deletion of *Lrh1* results in minimal impact on the gross crypt structure and the number of tdTomato-labeled cells per crypt. Scale bar is 50µm. N =3 per condition, data are presented as mean ± SEM. Statistical significance was determined using unpaired two-tailed Student’s t-test.

### *Lrh1* is dispensable in Atoh1^+^ cells for lineage maintenance in healthy tissue

Previous studies have revealed a supportive role for LRH-1 in the intestinal epithelium^4,5^. To mechanistically investigate how LRH-1 impacts intestinal homeostasis and repair, we utilized a mifepristone (RU486)-inducible Atoh1^CrePGR^ to selectively delete the *Lrh1* gene and activate the reporter allele tdTomato (tdT) in Atoh1^+^ cells of the secretory lineage, but not in ISCs^20^ (Fig. 1D). Following administration of RU486 to either mice via intraperitoneal (IP) injection or intestinal organoids, we observed reporter expression in intestinal crypts (Fig. 1D). Atoh1 expression of the FACS-sorted tdT-labeled cells was confirmed by RT-qPCR and RNAScope on the intestinal epithelium (Suppl. 1A-1C). *Lrh1* knockout was similarly confirmed in tdT^+^ sorted cells by RT-qPCR (Suppl 1D).

Because prior studies using the broadly expressed Vil^Cre-ERT2^ recombinase demonstrated that acute loss of *Lrh1* triggered cell death in intestinal organoids and compromised intestinal epithelial function in mice^4^, we asked whether deletion of *Lrh1* in Atoh1^+^ cells similarly compromised cell homeostasis. Examination of intestinal crypts following 3-day treatment with RU486 revealed a comparable proportion of tdT^+^ crypt cells along with preserved crypt architecture, suggesting that acute *Lrh1* loss in this population did not trigger significant cell loss or structural disruption (Fig. 1E). Furthermore, loss of *Lrh1* in Atoh1^+^ cells did not alter the number of ISCs as marked by *Olfm4* expression (Suppl. 1E).

### *Lrh1* is critical for Atoh1^+^ lineage-mediated ISC regeneration after radiation injury

Because *Lrh1* has been shown to have an important role in ameliorating intestinal epithelial damage^4^, we next explored whether it contributes to ISC regeneration as part of the intestinal healing process. We utilized the radiation enteritis model where mice are exposed to 8Gy total body irradiation (TBI) to cause small intestinal injury and ISC loss^21^. This radiation enteritis model has been widely used to investigate the dedifferentiation of committed intestinal epithelial cells and subsequent repopulation of ISCs^19,22-24^. Intestinal epithelial damage commences during the first 2 days following radiation, with sublethal damage repaired by one-week post-radiation with normal appearing histology and reappearance of ISCs^21^. To induce Lrh1 knockout and enable lineage tracing of Atoh1^+^ cells, mice received RU486 IP for three successive days prior to TBI (Fig. 2A). Intestinal tissues were harvested 10 days after TBI (Fig. 2B) to assess the impact of LRH-1 on ISC restitution and epithelial healing. Unlike other intestinal damage models utilizing haploinsufficiency or Vil^Cre^ drivers, selective *Lrh1* deletion in Atoh1^+^ cells did not contribute to exacerbated disease severity as measured by body weight and survival (Suppl. 2A, 2B), likely owning to the relatively small portion of epithelial cells undergoing *Lrh1* knockout in this model^4,5^. Remarkably, however, LRH-1 deletion resulted in a >50% reduction of new ISCs arising from Atoh1^+^ cells as determined by co-positivity of tdT and *Olfm4* in comparison with the LRH-1-sufficient control (Fig 2B,C).

**Figure 2.**
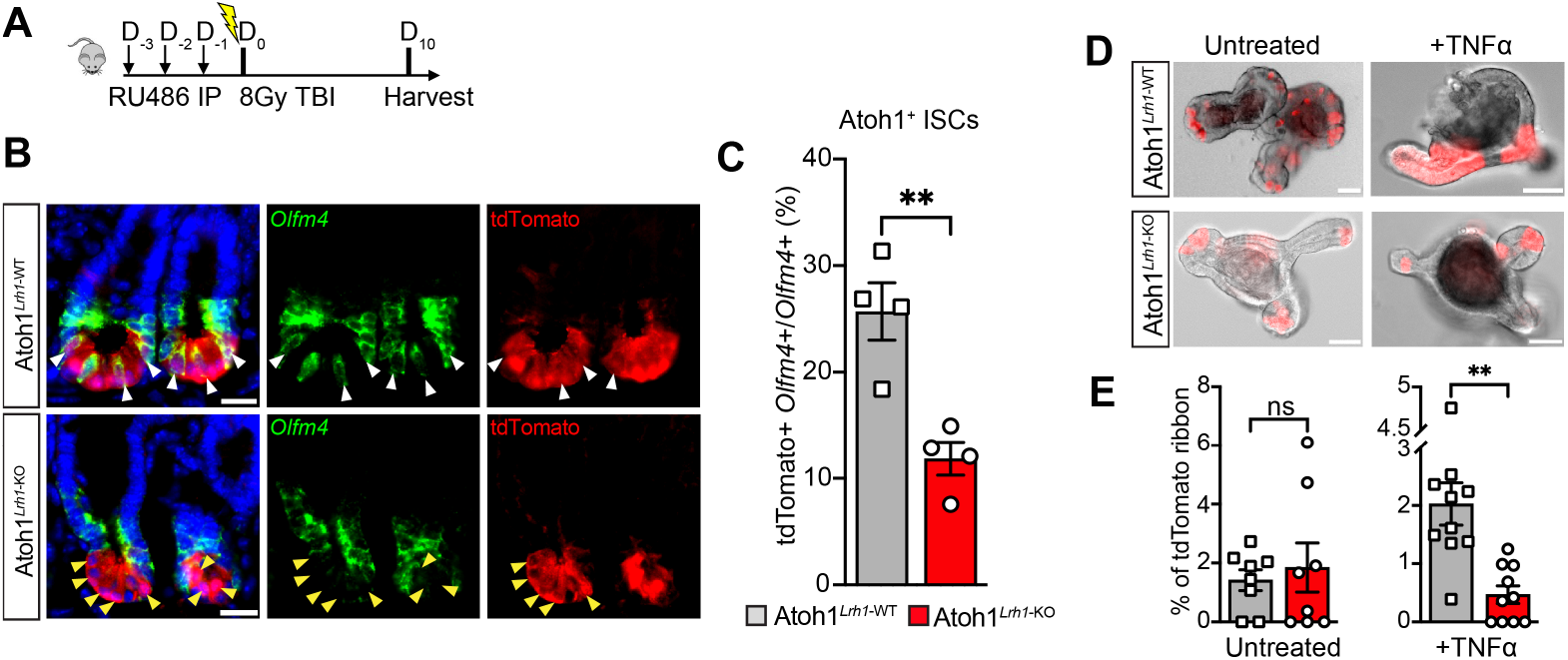
Lrh1 deletion in the Atoh1^+^ lineage impairs dedifferentiation after injury. **(A)** Schematic of radiation-induced enteritis model. **(B)** Crypts of *Lrh1*^Atoh1-WT^ (upper) and *Lrh1*^Atoh1-KO^ animals (lower) 10 days after radiation. New ISCs marked by co-expression of *Olfm4* and tdT^+^ (white arrowheads) seen predominantly in *Lrh1*^Atoh1-WT^ but not *Lrh1*^Atoh1-KO^. tdT^+^ cells lacking *Olfm4* expression in *Lrh1*^Atoh1-KO^ animals are predominantly non-ISC Paneth cells (yellow arrowheads). Scale bar is 20 µm. **(C)** Quantification of *Olfm4*^*+*^, tdT^+^ ISCs from dedifferentiated pre-labeled Atoh1^+^ cells in *Lrh1*^Atoh1-WT^ and *Lrh1*^Atoh1-KO^ animals. N=4 per condition. Small intestinal organoids derived from *Lrh1*^Atoh1-WT^ (upper) and *Lrh1*^Atoh1-KO^ (lower) mice show decreased lineage tracing as marked by tdT ribbon formation after *ex vivo* TNFα challenge. Scale bar is 50µm. **(E)** Quantification of tdT ribbon-positive organoids after TNFα treatment shows impairment of ISC repopulation from Atoh1^+^ cells following LRH-1 loss. n > 6 independent batches from at least 3 animals per condition.

To determine whether the LRH-1 impact on ISC repopulation is epithelial cell intrinsic or the result of a more complex interplay between the luminal environment or subepithelial interactions with mesenchymal or inflammatory cells^25-27^, we utilized intestinal organoids derived from *Lrh1*^*Atoh1-KO*^ and control animals. tdT-labeled Atoh1^+^ lineages were found in both crypt and villus compartments, mostly with a scattered pattern under basal homeostasis (Fig. 2D). Lineage-labeled Atoh1^+^ intestinal epithelial cells retain ISC properties and can form a clonal ribbon marked by tdT positivity spanning across the crypt-villus domains under homeostatic conditions^20^. Consistent with our in vivo observation of healthy intestine (Suppl. 2A), the percentage of organoids exhibiting tdT-labeled ribbons derived from Atoh1^+^ IECs remained unaffected by RU486-mediated Lrh1 knockout under baseline conditions (Fig. 2D,E, untreated). To mimic injury-associated inflammation ex vivo, organoid cultures were challenged with Tumor Necrosis Factor-alpha (TNFα; 10 ng/ml for 40 hours)^4,28-31^. Organoid cultures were maintained for an additional 48h after removal of TNFα prior to imaging analysis to enable assessment of reparative dedifferentiation and clonal expansion. As in our radiation enteritis model, *Lrh1* loss resulted in a greater than 50% reduction in clonal ribbon formation as compared with LRH-1 sufficient controls (Fig. 2D,E).

Taken together, these results using complementary in vivo and ex vivo intestinal injury models identify a vital epithelial-intrinsic role for Lrh1 in promoting dedifferentiation of the Atoh1^+^ lineage and ISC restitution following injury.

### Transcriptomic analysis implicates impaired protein processing in the endoplasmic reticulum after LRH-1 deletion

To explore the mechanisms underlying impaired ISC regeneration associated with LRH-1 loss, we performed bulk RNA sequencing on FACS-sorted tdT^+^ cells isolated from *Lrh1*^*Atoh1-WT*^ and *Lrh1*^*Atoh1-KO*^ intestines of 8-to 10-week-old mice (Fig. 3A). Density distribution of FPKM and principal component analysis (PCA) of the ∼24,000 detected genes showed similar transcriptomes (Suppl. 3A, 3B), consistent with the grossly normal crypt-villus architecture in unperturbed *Lrh1*^*Atoh1-KO*^ intestinal epithelium. However, gene set enrichment analysis (GSEA) uncovered alterations in endoplasmic reticulum (ER)-associated pathways and protein processing (Fig. 3B-D). Independent GO, KEGG, and Reactome enrichment analysis confirmed downregulation of ER-associated processes in *Lrh1*^*Atoh1-KO*^ cells (Fig. 3B-3D). GO terms related to ER protein processing were enriched across molecular functions, cellular components, and biological process categories. KEGG pathway analysis pinpointed protein processing in ER (MMU04141) as a significantly compromised pathway following *Lrh1* knockout. Reactome analysis further ranked the unfolded protein responses (UPRs) that resolve misfolded protein-induced ER stress as top-downregulated gene sets by adjusted p-value out of 159 identified downregulated genes. Collectively, these analyses suggest impairment in ER protein processing and UPRs in *Lrh1*^*Atoh1-KO*^ animals relative to *Lrh1*^*Atoh1-WT*^ controls.

**Figure 3.**
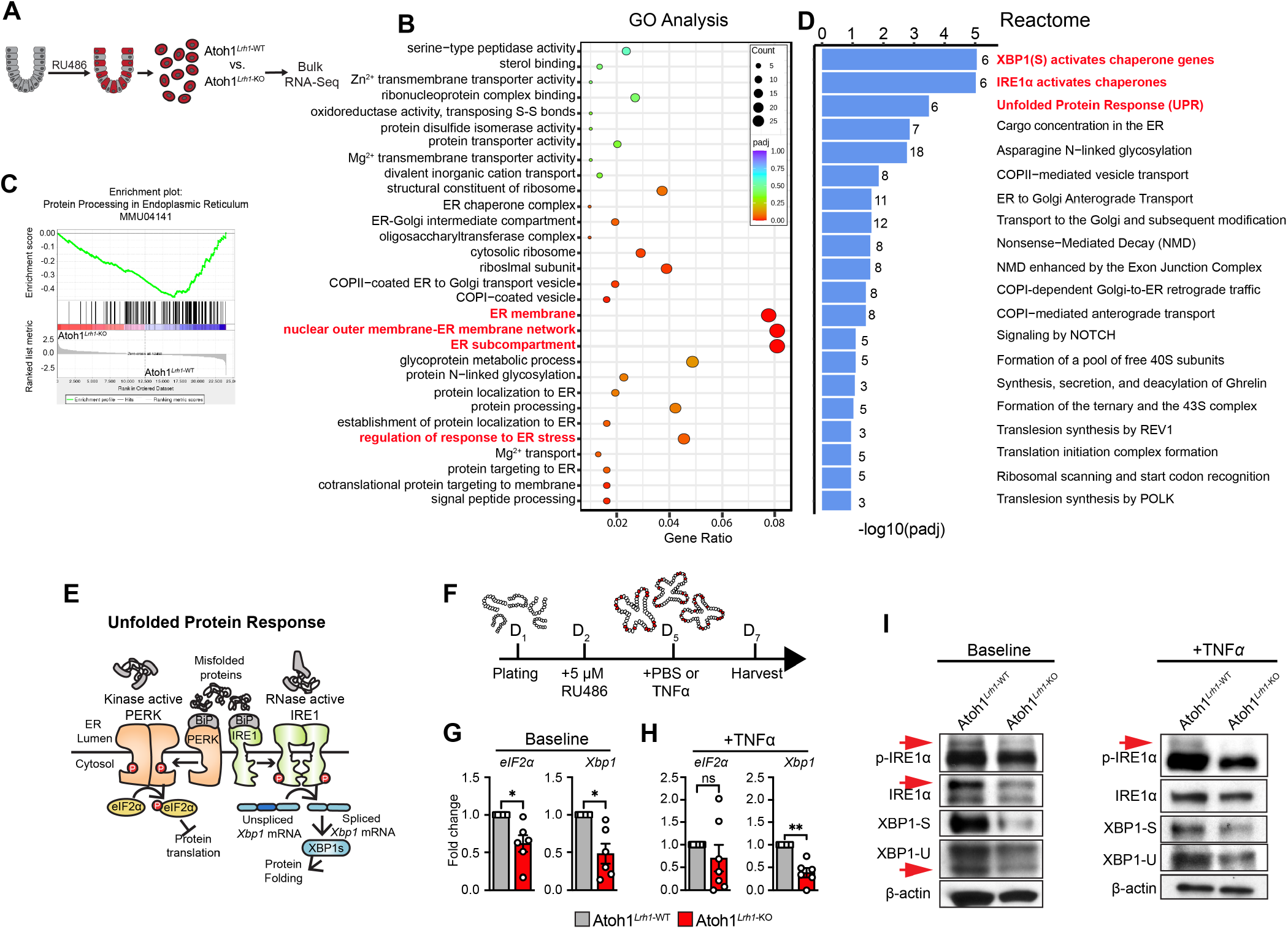
*Lrh1* knockout impairs the IRE1α-XBP1 Unfolded Protein Response. **(A)** Schematic diagram illustrating the workflow of the bulk RNA-seq. **(B)** Gene ontology (GO) analysis identifies that *Lrh1*^Atoh1-KO^ primarily affects gene functions in ER-related protein folding processes (red). **(C)** KEGG analysis suggests protein processing in the endoplasmic reticulum is impaired in *Lrh1*^Atoh1-KO^. **(D)** Reactome analysis finds *Lrh1*^Atoh1-KO^ predominantly impacts the IRE1α-XBP1 pathway. **(E)** Schematic illustration of PERK-eIF2α and IRE1α-XBP1 branches of the UPRs responsible for resolving misfolded protein-induced ER stress. Misfolded peptides trigger PERK and IRE1α dimerization and autophosphorylation causing PERK-eIF2α-mediated attenuation of protein translation and IRE1α-directed splicing of Xbp1 mRNA to promote UPR target gene expression. **(F)** Schematic illustrating the intestinal organoid TNFα challenge experiment. **(G)** RT-qPCR from unsorted organoids showed reduced gene expression of *eIF2α* and *Xbp1*, the executors of PERK and IRE1α branches of UPRs at steady state. **(H)** RT-qPCR revealed that expression of *Xbp1*, but not *eIF2α*, in unsorted *Lrh1*^Atoh1-KO^ organoids is significantly impaired after TNFα treatment. **(I)** Western blot of unsorted *Lrh1*^Atoh1-WT^ and *Lrh1*^Atoh1-KO^ organoids at baseline (left) and after TNF*α* treatment (right). *Lrh1*^Atoh1-KO^ led to reduction in total IRE1α, p-IREα, and XPB1 protein levels. Red arrow indicates bands of interest. For panels G and H, n=6 independent batches from 3 animals per condition. Data are presented as mean ± SEM. Statistical significance was determined using unpaired two-tailed Student’s t-test. * p < 0.05, ** p < 0.01

### *Lrh1* deletion in Atoh1^+^ lineage cells impairs the IRE1α-XBP1 unfolded protein response

Bulk RNA-seq revealed that Atoh1^+^ lineage of *Lrh1*^*Atoh1-KO*^ animals exhibited altered expression of transcripts involved in ER-associated protein processing and UPR, suggesting that *Lrh1* loss impairs the cellular ER stress response. The ER is the primary organelle for quality control of newly synthesized polypeptides where they undergo proper folding and modification before trafficking to their final destination. Accumulation of misfolded proteins in the ER triggers the ER stress response, which recruits the adaptive UPR to restore proteostasis for cell survival^32-34^ (Fig. 3E). To understand how LRH-1 modulates the ER stress response, we interrogated our RNA-seq data and identified multiple candidate genes including elements involved in importation of nascent polypeptide into the ER (*Sec61a, Sec62*), translocation of proteins across the ER membrane and regulating N-linked glycosylation (*Ssr1-4*), and core UPR components. Individual validation by RT-qPCR in unsorted organoids derived from *Lrh1*^*Atoh1-KO*^ mice confirmed loss of LRH-1 reduced expression of *eukaryotic translation initiation factor 2, subunit 1 alpha* (*Eif2s1*, also known as *eIF2α*) and *X-box binding protein 1* (*Xbp1*), both critical components mediating the UPR response (Fig. 3F,G), but not *Sec61a, Sec62*, or *Ssr1-4* (Suppl. 4a).

To assess LRH-1 function in an inflammatory context, we challenged intestinal organoids with TNFα (20ng/ml for 24 hours), a stimulus known to induce ER stress^35^. RT-qPCR analysis showed expression of *efF2α* remained statistically unchanged in *Lrh1*^*Atoh1-KO*^ organoids relative to *Lrh1*^*Atoh1-WT*^ controls. In contrast, *Xbp1* expression was significantly reduced in TNFα-challenged unsorted *Lrh1*^*Atoh1-KO*^ organoids (Fig. 3H). Under stress conditions, IRE1α undergoes phosphorylation and dimerization, activating it to splice *Xbp1* mRNA into its active form, *Xbp1s*. The resulting XBP1s protein translocates to the nucleus and activates UPR target genes to alleviate ER stress (Fig. 3E). Western blot analysis of unsorted organoids revealed that LRH-1 deletion in Atoh1^+^ lineage cells reduced total IRE1α and phosphorylated IRE1α under basal and TNFα-challenged conditions (Fig. 3I). Similarly, protein levels of unspliced XBP1 (XBP1u) and spliced XBP1 (XBP1s) were decreased in unsorted *Lrh1*^*Atoh1-KO*^ organoids regardless of TNFα treatment (Suppl. 4B). Taken together, these findings suggest that LRH-1 supports the IRE1α-XBP1 branch of UPR in the intestine, and that its loss impairs a compensatory cellular response to injury-induced ER stress.

### Ire1α-Xbp1 are candidate gene targets for LRH-1

Knockout of *Lrh1* in the Atoh1^+^ lineage resulted in reduced RNA expression and protein levels of IRE1α and XBP1 (Fig. 3), suggesting these genes might be direct transcriptional targets of LRH-1. Bioinformatic analysis of the 2kb promoter regions upstream of their respective transcriptional start sites using the Eukaryotic Promoter Database software program^36,37^ identified multiple putative LRH-1 binding sites (Fig. 4A). Because of the unsuitability of mouse LRH-1-directed antibodies for immunoprecipitation and downstream biochemical analyses, we utilized the CRISPR/Cas9 platform to knock in a 3xFLAG tag into the endogenous Lrh1 locus at the C-terminal of exon 9 (Fig. 4B). Successful knock-in was confirmed by genotyping PCR (Fig. 4C) and anti-FLAG western blot from intestinal tissue and organoid lysate (Fig. 4D). The resulting *Lrh1-3xFLAG* line was backcrossed for seven generations to the C57BL/6J background to minimize genetic background effects. Using this new LRH-1 tool, we performed chromatin immunoprecipitation followed by quantitative PCR (ChIP-qPCR) assays on isolated intestinal crypts for *Ire1α* and *Xbp1* promoter regions. Compared with IgG control, ChIP-qPCR derived from anti-FLAG lysates demonstrated significant enrichment at the predicted LRH-1 binding sites within the promoter regions of both genes (Fig. 4E). These results provide evidence that LRH-1 directly binds to the promoters of *Ire1α* and *Xbp1* and supports a potential direct role in regulating their transcription.

**Figure 4.**
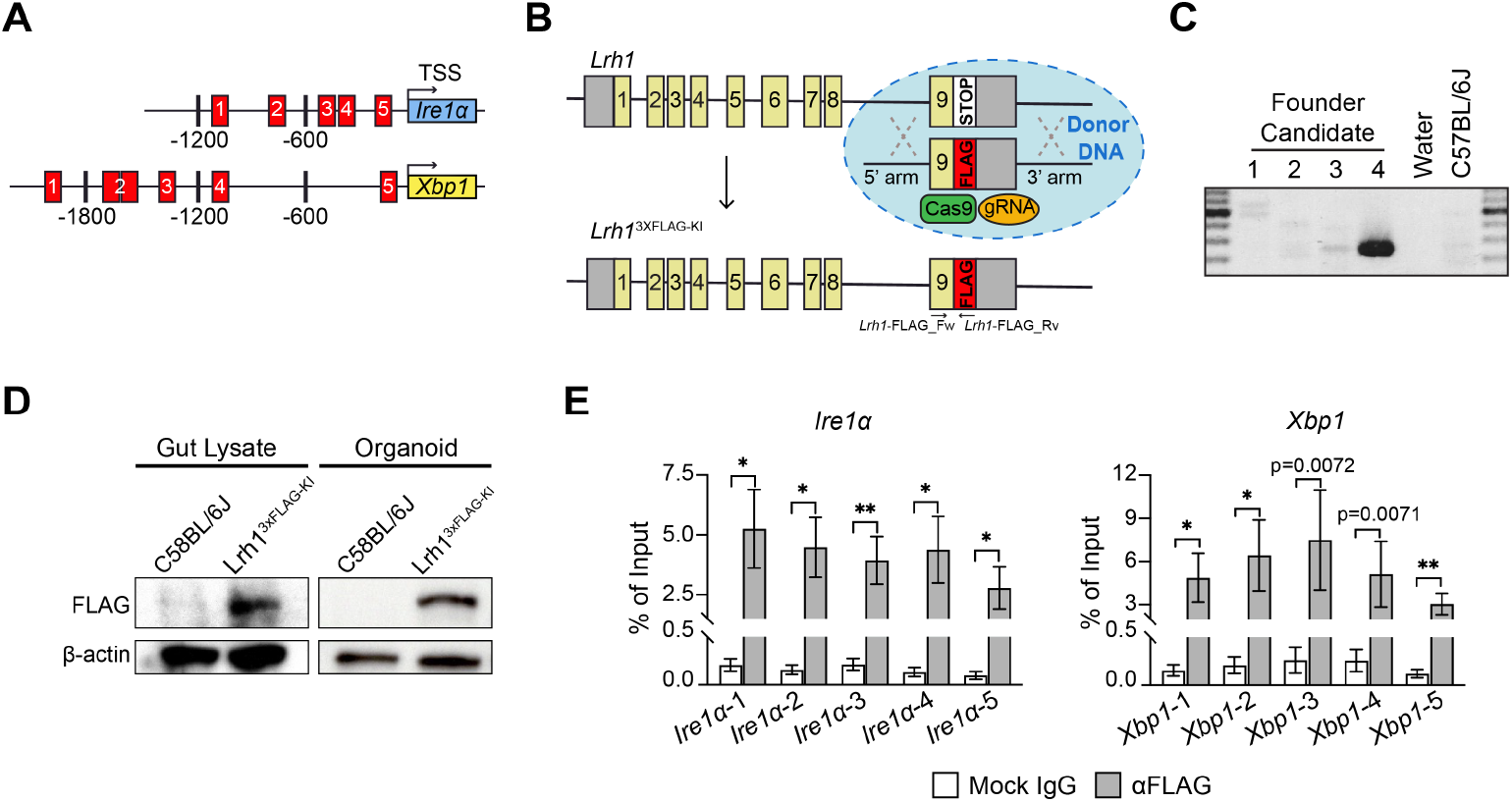
LRH-1 binds to the promoter regions of *Ire1α* and *Xbp1* genes. **(A)** Schematic diagram illustrating the predicted LRH-1 binding sites within the 2000 bp upstream to the transcription starting site (TSS) of *Ire1α* and *Xbp1*. **(B)** Schematic diagram depicts the generation of *Lrh1*-3XFLAG knock-in mouse. C57BL/6J mouse *Lrh1* genomic locus (top) and the CRISPR-mediated insertion of the DNA sequence of a 3XFLAG epitope tag at the C terminus of *Lrh1* locus(bottom). **(C)** PCR validation of genomic targeted knock-in of the 3XFLAG in F_0_ founders. **(D)** Western blot assay validates the protein expression of 3xFLAG in mouse intestine and intestinal organoid as detected by FLAG antibody. **(E)** ChIP-qPCR results from adult *Lrh1*-3XFLAG knock-in mouse intestine indicate LRH-1 binding at *Ire1α* and *Xbp1* promoter region. N=8. Data are presented as mean ± SEM. Statistical significance was determined using unpaired two-tailed Student’s t-test. * p < 0.05, ** p < 0.01

## Discussion

In this study, we employed complementary in vivo and ex vivo intestinal injury models to establish that LRH-1 plays a critical role in post-injury intestinal epithelial regeneration. For the first time, we showed that acute LRH-1 deletion in Atoh1^+^ cells impaired the capability of Atoh1-expressing intestinal epithelial cells to dedifferentiate and repopulate the crypt base ISC pool after tissue damage. We further revealed that loss of LRH-1 impaired the expression of key cellular injury genes upon exposure to TNFα, a major inflammatory cytokine elevated in IBD. Transcriptomic and protein level analyses of LRH-1-deficient Atoh1^+^ intestinal epithelial cells indicate that LRH-1 engages the IRE1α-XBP1 branch of the unfolded protein response. Using our newly developed *Lrh-1-3xFLAG* knock-in mouse to perform ChIP-qPCR, we found enriched LRH-1 binding at predicted promoter binding sites of *Ire1α* and *Xbp1* genes in native intestinal tissue, providing evidence for LRH-1-mediated transcriptional regulation of this critical stress-responsive pathway. Taken together, our findings complement prior studies demonstrating an important role for LRH-1 in ameliorating colitis and provide a new mechanism whereby this nuclear receptor supports intestinal epithelial repair via progenitor cell dedifferentiation and ISC restitution.

Our results demonstrate that LRH-1 is critical for ISC restitution following intestinal epithelial injury. By using selective, inducible LRH-1 deletion confined to a subset of non-ISC intestinal epithelial cells, we uncovered a new epithelial cell-intrinsic role for LRH-1 in injury-induced regeneration not detectable in previous studies utilizing the broadly expressed Cre drivers^4,5^. In both ex vivo intestinal organoids challenged with TNFα and in vivo radiation-induced enteritis, we found that LRH-1 deficiency substantially reduced the generation of new ISCs from dedifferentiated Atoh1^+^ cells. Atoh1 marks secretory progenitor cells, which are known to undergo dedifferentiation upon tissue injury, and its expression is persistent in differentiated goblet and Paneth cells, which can also contribute to ISC repopulation in damage contexts^15,22^. In our study, we are unable to distinguish between these Atoh1^+^ secretory lineage sources. Another limitation of our model is incomplete recombination in all Atoh1-expressing cells, with an apparent bias towards Paneth cells as evidenced by co-positivity of the tdTomato reporter and Paneth cell marker lysozyme in a high proportion of labeled crypt-base cells (Suppl. 5A). This restricted labeling efficiency likely explains the relatively lower number of reporter-labeled newly formed ISCs in our control group compared with previous secretory progenitor reports^38^. Nevertheless, our findings extend the known protective roles of LRH-1 in the intestinal epithelium and reveal a novel mechanism whereby LRH-1 promotes epithelial repair and ISC recovery after injury.

Through integrated gene expression pathway analysis and biochemical assessment of key regulatory proteins, we provide evidence that LRH-1 promotes the UPR to mitigate ER stress, protecting intestinal epithelial cells from acute cellular injury. Our findings are in line with previous reports of LRH-1 modulating ER stress in a hepatic injury model^39^. In the intestine, we found that LRH-1 selectively engages the IRE1α-XBP1 branch of the UPRs, as evidenced by reduced transcript and protein levels of Ire1α and Xbp1 in *Lrh1* knockout cells. Using our newly constructed *Lrh1-3xFLAG* knock-in mouse, we confirmed direct binding of LRH-1 to the promoter regions of both *Ire1α* and *Xbp1*, strongly supporting a role for LRH-1 in transcriptional regulation of these key genes as part of an injury-adaptive program. In contrast, hepatic LRH-1 alleviates cellular injury primarily through *polo-like kinase 3* (*Plk3*) to resolve chemical-elicited ER stress. Interestingly, activation of the PERK-eIF2α arm of the UPR has been linked to compromised stemness in various intestinal contexts^40,41^. Our results suggest two plausible explanations: (1) the IRE1α-XBP1 branch functions independently of PERK-eIF2α branch in preserving ISC properties during repair, or (2) LRH-1-dependent UPR enhances the resilience of Atoh1^+^ lineage cells under stress, thereby promoting cell survival and enabling subsequent dedifferentiation and ISC restitution. These branch-specific UPRs underscore the context-dependent distinction in maintaining intestinal tissue homeostasis.

Prior literature has established pro-survival and anti-inflammatory roles for LRH-1 across multiple organs, including the intestine. Interestingly, complete or partial loss of LRH-1 appears to be relatively well tolerated under homeostatic conditions^4,42,43^. However, upon tissue injury LRH-1 insufficiency leads to exaggerated inflammatory damage and impaired subsequent repair^4,5^. Our findings with inducible Lrh1 deletion mediated by Atoh1^CrePGR^ recapitulate and extend these observations. At baseline, loss of LRH-1 in Atoh1^+^ cells did not cause detectable alterations in crypt-villus structure or in the maintenance of tdTomato-labeled Atoh1^+^ lineage cells. In contrast, both ex vivo organoids challenged with TNFα and in vivo radiation-induced tissue injury revealed clear defects associate with LRH-1 loss, including compromised cellular stress responses and ISC regeneration. Given that our model targets a relatively small proportion of intestinal epithelial cells, subtle homeostatic defects in LRH-1 deficient cells may exist but elude detection in the absence of tissue injury.

An unresolved question is whether LRH-1 exerts part of its protective activity through Paneth cells, specialized secretory cells that play critical roles in antimicrobial defense and in support of the ISC niche. LRH-1 expression appears to occur at low levels in a subset of Paneth cells as indicated by RNAscope co-localization of Lrh1 and the Paneth cell marker lysozyme (Suppl. 5B). Given the dependence of highly secretory, metabolically active cells on ER stress and UPR pathways for cellular homeostatic maintenance and antimicrobial peptide production, LRH-1 may contribute to maintaining Paneth cell health and viability via the IRE1α-XBP1 pathway. Considering the robust Lrh1 expression in the progenitor compartment (Fig. 1A), an alternative explanation is that LRH-1 loss in these transient amplifying cells, including Paneth cell progenitors, impairs proper maturation and the subsequent niche-supporting functions of mature Paneth cells. Dissecting the relative contributions of LRH-1 effects in mature Paneth cells versus progenitor defects will be the focus of future study.

A major deficit in the LRH-1 field has been a lack of reliable antibodies to enable targeted in vivo and ex vivo molecular analyses. Here we show the suitability of our novel LRH-1-FLAG3x construct for performing LRH-1-chromatin analysis from native tissue as proof-of-construct. We expect LRH-1-FLAG3x will enable new precision molecular studies broadly across tissues types and disease models including pancreas^42^ and liver^39^.

Our findings provide strong support for therapeutically targeting LRH-1 in intestinal disorders. Previous studies have established that overexpression of LRH-1 in the intestinal epithelium confers resistance to colitis in an adoptive immune transfer model^4,44^. More recently, preclinical pharmacological agonists of LRH-1 tested in mice humanized for intestinal LRH-1 expression showed efficacy in ameliorating experimental colitis^6^.

Taken together with our results, these observations suggest that targeted activation of LRH-1 may provide a therapeutic benefit in inflammatory conditions such as IBD, graft-versus-host disease, and checkpoint inhibitor-induced colitis via multiple complementary actions. Pharmacological enhancement of LRH-1 activity could promote cell survival by augmenting the IRE1α-XBP1 branch of the unfolded protein response, facilitating ISC regeneration and niche restoration after injury, and activation of anti-inflammatory and immunoregulatory gene programs^45,46^. Although further studies are needed to assess the safety, specificity, and efficacy of LRH-1 agonists in relevant humanized animal model or human systems, our report positions LRH-1 as a promising target for drug design strategies that aim at enhancing epithelial repair and mucosal healing in chronic inflammatory intestinal diseases.

## Acknowledgements

We thank Drs. Holly Ingraham, Shaoyi Zhang, Shuo Wang and all members of the Bayrer laboratory for helpful discussions. We thank S.A. Kliewer (UT Southwestern Medical Center) for the gift of Lrh1f/f mice. We additionally thank Drs. Diego Miranda, Candice Herber for helpful discussions in the early stages of this project. This work was supported by R01DK128346 (JRB), R03DK121061 (JRB), UCSF Chancelor’s Fund and Chan Future of UCSF (JRB), P30DK098722 Pilot & Feasibility Award (JRB), and T32DK007762 (HJC).

## Materials and Methods

### Animals

All mouse experiments were approved by the Institutional Animal Care and Use Committee (IACUC) of the University of California, San Francisco and were performed with the guidelines. *Atoh1*^CrePGR^, Rosa26-LSL-tdTomato, and *Lrh1*^flox/flox^ mice were described previously^47^, ^48^, ^49^. *Atoh1*^*CrePGR*^ was a gift from Huda Y. Zoghbi, M.D. (Baylor College of Medicine). All experiments were performed on 10-14 weeks old animals, both male and females were randomly used for all experiments. Mice were housed in a specific pathogen-free, temperature-controlled room with 12-hour light-dark cycle and were fed the normal diet food and water. To induce Cre-mediated recombination, mifepristone (RU486; 2mg/body) was administrated intraperitoneally into mice that carry the *Atoh1*^*CrePGR*^ allele for 3 successive days^20^.

### Intestinal epithelial organoid culture

Isolation and culture of the primary intestine crypts into 3-dimension enteroid was carried out as previously described^50^, ^51^. Briefly, the entire small intestine was splayed open and rinsed with cold 1x PBS. After gently scraping with a glass coverslip to remove the villi and mucus, tissue was sliced into 2 mm-width pieces and was incubated in cold 1x PBS containing 5mM EDTA at 4 °C for 45 minutes. Followed by vigorous agitation, the crypt-enriched suspension was filtered through 70-µm cell strainer and then centrifuge at 300 ×g, 4°C for 5 minutes, washed in cold 1x PBS, and pelleted again at 300 ×g, 4°C for 5 minutes. Crypts were resuspended in Matrigel droplets (Corning, 354234) and cultured in the complete ENR medium that consists of advanced DMEM/F12 medium (Thermo Fisher Scientific, 12634010), 1x GlutaMax (Thermo Fisher Scientific, 35050061), 10 mM HEPES (Thermo Fisher Scientific, 15630080), 1x Penicillin-Streptomycin (Thermo Fisher Scientific, 15140122), 1x N-2 Supplement (Thermo Fisher Scientific, 17502048), 1x B-27 Supplement (Thermo Fisher Scientific, 17504044), 1 mM N-Acetyl-l-cysteine, 50 ng/mL mEGF (Thermo Fisher Scientific, PMG8041), 100 ng/mL mouse Noggin (Peprotech, 250-38-500UG), 10% R-spondin conditioned medium (Kind gift from Jeffrey Whitsett, Cincinnati Children’s Hospital Medical Center). To induce the *Atoh1*^CrePGR^-mediated lineage labelling *ex vivo*, 5uM RU486 in 100% ethanol was added in the complete ENR medium 24 hours before further treatments.

### RNAscope fluorescent in situ hybridization

RNAscope fluorescent *in situ* hybridization was performed on OCT-embedded tissue section according to RNAscope multiplex fluorescent reagent Kit v2 manual with adaptation. Briefly, cryosections were baked in the oven at 60°C for 30 minutes prior to incubation in 4% PFA at room temperature (RT) for 30 min. Slides were dehydrated with 5-minute wash in 50%, 70%, and two successive washes in 100% ethanol at RT. After drying at 60°C for 10 minutes, slides were incubated with hydrogen peroxide (H_2_O_2_) at RT for 10 minutes to inactivate endogenous peroxidase activity and reduce non-specific background. For target retrieval, slides were heated in a steamer at >95°C in 1x target retrieval solution (Advanced Cell Diagnostics, 322000) for 5 minutes, rinsed with fresh water, dehydrated in 100% ethanol, and treated with Protease III at 40°C for 30 minutes. Probe hybridization was performed with target-specific probes at 40°C for 2 hours. Signal amplification was conducted using RNAscope Multiplex Fluorescent Reagent Kit v2 (Advanced Cell Diagnostics, 323100), followed by tyramide signal amplification (TSA) with Opal® fluorescent dyes (1:1500, Perkin Elmer, FP1487A). For slides co-immunostaining after RNAscope *in situ* hybridization, samples were blocked in 5% normal serum/0.1% Triton X-100 in 1x PBS at RT for 1 hour and then incubated overnight at 4°C with primary antibodies. Following three washes in 0.1% Triton X-100 in 1X PBS, slides were incubated with Alexa Fluor-conjugated secondary antibodies (1:1000). Nuclei were counterstained with DAPI (1 µg/ml).

### Immunostaining

Mouse tissue was fixed in 4% paraformaldehyde at 4°C for overnight, followed by 30% sucrose solution incubation at 4°C for overnight before embedding in OCT-medium. To unmask the antibody-binding epitopes, 5µm OCT-medium embedded sections were soaked in the 95°C preheated antigen retrieval buffer (1uM citric acid, pH 6.0) for 15 min and then transferred into 1x PBS to remove the retrieval buffer. After blocking with 5% normal serum in 0.1% Triton-X100 in 1x PBS (PBS-T) at RT for 1 hour, slices were stained with assigned primary antibodies at 4°C for overnight, followed by PBS-T washing and incubation with secondary antibodies at RT for 1 hour. Nuclei were counterstained with DAPI (1 µg/ml) and mounted with Fluoromount-G™ Mounting Medium (Invitrogen, 00-4958-02). All stained slices were imaged on the Keyence BZ-X1000 or Crest LFOV Spinning Disk/ C2 Confocal microscope.

### Whole-body irradiation

Animals were injected with RU486 intraperitoneally for 3 consecutive days before receiving 8 Gy whole-body irradiation (Precision X-RAD 320 Cabinet Irradiator, Caseium-137). To ensure the body condition score complied with the UCSF animal welfare policy, all post-irradiation animals were individually housed with supportive heating pads and received daily body weight measurement.

### Fluorescence-Activated Cell Sorting

To label the *Atoh1*-positive lineages *in vivo, Lrh1*^WT/WT^; *Atoh1*^CrePGR^, Rosa26-LSL-tdTomato (*Lrh1*^Atoh1-WT^) and *Lrh1*^flox/flox^; *Atoh1*^CrePGR^, Rosa26-LSL-tdTomato (*Lrh1*^Atoh1-KO^) was given RU486 (2mg/body) intraperitoneally 3 consecutive days prior to tissue harvest. Single cell dissociation was performed as previous described^22^. After washing the isolated crypts with cold 1x PBS, the pellets were resuspended with TrypLE Express (Thermo Fisher Scientific, 12604013) at 37 °C for 3 minutes. The cell suspensions were mixed with 20% FBS in 1x PBS to quench the TrypLE-mediated dissociation, filtered through a 40µm strainer, and centrifuged at 300 ×g, 4 °C for 5 minutes. Cell pellet was washed with cold 1x PBS and resuspended in FACS sample buffer (DMEM without phenol red containing 2% FBS and 50 unit/ml DNase I). Cells were incubated with Alexa Fluor™ 700 - conjugated CD45(1:200) and FITC-conjugated EpCAM (1: 200) on ice for 30 minutes. DAPI was mixed immediately before sorting to label alive cells. FACS was performed with BD Biosciences Aria II Flow Cytometer. Single viable epithelial cells were gated by negative staining of DAPI and CD45, and positive staining of EpCAM. *Atoh1*-positive population was further collected based on RU486-mediated expression of tdTomato. FACS data were analyzed using FlowJo™ (v11.1) Software. Viable tdTomato-positive singlets were directly sorted into the RLT lysis buffer (Qiagen 79216) supplemented with 1% β-mercaptoethanol and stored in -80°C for the following bulk RNA-Seq analysis.

### Bulk RNA-Seq Analysis

RNA of sorted *Atoh1*^+^ lineage from *Lrh1*^Atoh1-WT^ and *Lrh1*^Atoh1-KO^ were extracted with PicoPure™ RNA Isolation Kit (Applied Biosystems™, KIT0204), eluted in 12 µl RNase-free water, and sequenced using SMART-Seq® v4 Ultra® Low Input RNA (Takara, 634888). RNA reads were aligned with HISAT2 (2.0.5) and visualized using Integrative Genomics Viewer (2.15.10). Aligned read files were run simultaneously using FeatureCount (1.5.0-p3) to quantify the values of Fragments Per Kilobase of transcript per Million mapped reads (FPKM) with default setting. For differential gene expression analysis, FPKM-based statistical analysis of the expression was normalized, and genes were ranked using DESeq2 (1.20.0). ClusterProfiler (3.8.1) was used for enrichment analysis. Ranked gene list was further analyzed using GSEA (3.0).

### Quantitative Real time-PCR

Intestinal organoids were released from Matrigel droplets by incubating with cold 1x PBS and trituration. RNA was extracted using the QIAGEN RNeasy mini kit (Qiagen 74104), and cDNA was prepared with iScript™ cDNA Synthesis Kit (Bio-Rad, 1708891BUN). PCR reaction mixture was assembled by the PerfeCTa SYBR® Green SuperMix (Quantabio, 95053500). Each dataset was generated from at least three independent biological replicates. Each reaction was performed in technical triplicates by Bio-Rad CFX384 Touch Real-Time PCR Detection System. Parallel PCR using β-2-microglobulin (*B2m*) was the reference, and the expression of each gene was normalized to that of the *B2m* and set as 1 in the *Lrh1*^Atoh1-WT^ control sample. The expression of each gene in the *Lrh1*^Atoh1-KO^ was normalized to that of *B2m* and then calculated as fold change compared to that in the *Lrh1*^Atoh1-WT^ control group.

### Western blotting

Intestinal organoids were harvested from Matrigel droplets with cold 1x PBS and trituration. Cell pellets were lysed in 1x lysis buffer (62.5 mM Tris-HCL pH 6.7, 1% SDS, 10% Glycerol) supplemented with 1x Halt™ Protease Inhibitor Cocktail (Thermo Fisher Scientific, 78425), and protein concentration was measure by the Pierce BCA Protein Assay Kit (Thermo Fisher Scientific, 23227). Lysate samples were boiled for 10 minutes with 1X Laemmli Sample Buffer (Bio-Rad, 1610747) containing 2.5% β-mercaptoethanol and run on Mini-PROTEAN TGX™ Precast Protein Gels (Bio-Rad, 4561084) in 1x Tris-Glycine-SDS buffer (Bio-Rad, 1610732). Gels were transferred onto the PVDF membrane in 1x Tris-Glycine buffer (Bio-Rad, 1610734) containing 20% methanol. After blocking with 3% milk in TBST (0.1% Tween-20 in 1x TBS), PVDF membranes were incubated overnight at 4 °C with diluted primary antibody in 5% BSA/TBST, using the dilution recommended in the antibody data sheet. Blots were washed in TBST and placed with HRP-conjugated secondary antibody (1: 5000) at RT for 1.5 hours. Blot were incubated with SuperSignal™ West Atto Ultimate Sensitivity Substrate (Thermo Fisher Scientific, A38554) and imaged using X-ray films.

### Chromatin immunoprecipitation

Cell preparation and fixation were performed as previously described51, 52 with minor modification. Briefly, isolated crypts cells were pelleted and resuspended in DMEM with 1% methanol-free formaldehyde (Thermo Fisher Scientific, 28906) at RT for 15 minutes, followed by blocking in 125 mM glycine to quench the crosslink at RT for another 10 minutes. Cells were washed with pre-cold 1x PBS and pelleted at 500 ×g, 4 °C for 10 minutes. For nuclei isolation, the cell pellet was resuspended in 1 ml pre-cold Farnham Lab (FL) buffer (5mM PIPES pH 8, 85 mM KCl, 0.5% NP-40) supplemented with EDTA-free protease inhibitor (Thermo Fisher Scientific A32955) and sonicated in 1 ml AFA milliTUBE (Covaris 520135) using Covaris S220 Focused-ultrasonicator (peak power 75 W, duty factor 2%, 200 cycles per burst) at 4 °C until > 70% of nuclei were individualized. Isolated nuclei were collected by centrifugation, resuspended in 1ml pre-cold shearing buffer (10 mM Tris-HCl pH 8, 0.1% SDS, 1 mM EDTA) with EDTA-free protease inhibitor, and transferred to a new sonication tube. Chromatin was sheared using Covaris S220 (peak power of 50 W, a duty factor of 10%, 200 cycles per burst) at 4°C for 9 minutes to generate 200-500 bp fragments. Immunoprecipitation was performed overnight at 4°C with 5 µl anti-FLAG antibody (Sigma-Aldrich, F1804) or 5µl normal mouse IgG control (Santa Cruz Biotechnology, sc-2025) using chromatin from the crypt cells of Lrh1-3xFLAG knock-in mice. Antibody-bound chromatin was pulled down by Protein A/G Magnetic Beads (Thermos Fisher Scientific 26162), washed, and eluted. Following reversal of crosslink, DNA was purified by ChIP DNA Clean & Concentrator kit (Zymo Research D5205). Enrichment of target DNA sequence bound by LRH-1 was quantified by qPCR and expressed as the percentage of input DNA.

**Suppl 1.**
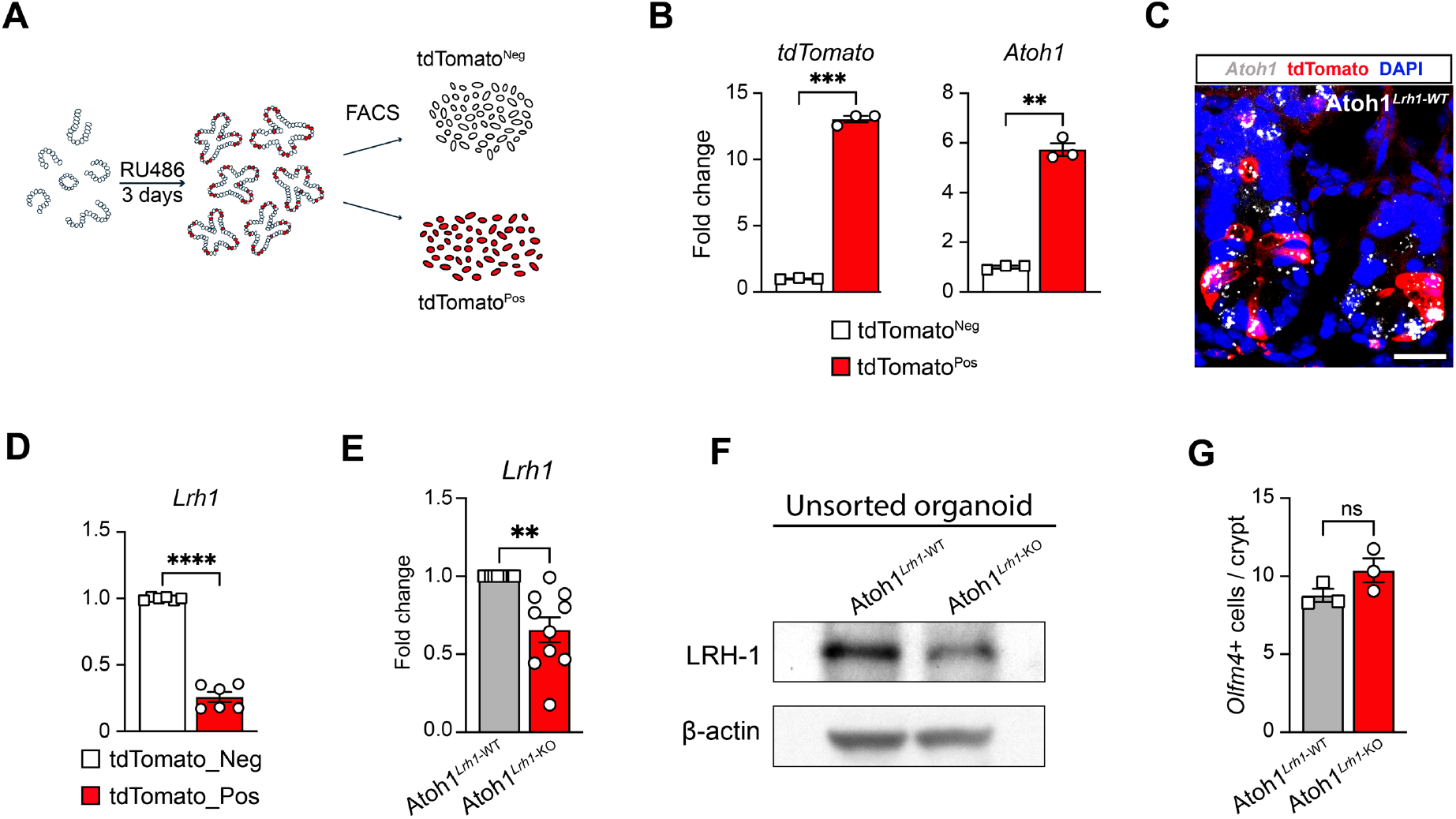
Confirmation of Atoh1^CrePGR^-mediated Atoh1^+^ lineage labeling. **(A)** Schematic diagram illustrating the workflow for validation studies. **(B)** RT-qPCR results from sorted organoids for *tdTomato* and *Atoh1*. **(C)** Representative image showing overlap of *Atoh1* (white) expression by RNAscope and tdT in mouse intestinal epithelium. Scale bar is 20 µm. **(D)** RT-qPCR result from sorted organoid confirmed reduced *Lrh1* expression in tdTomato-positive *Lrh1*^Atoh1-KO^ cells compared to its tdTomato-negative group. **(E)** RT-qPCR result from unsorted organoids showed reduced *Lrh1* expression following RU486 administration. **(F)** Western blot result from unsorted organoid showed reduced LRH-1 level under steady state. **(G)** Quantification of average number of *Olfm4*^+^ cells per crypt. *Lrh1* deletion by *Atoh1*^CrePGR^ results in minimal impact on the number of *Olfm4*^+^ cells per crypt. N=3 per condition. For panels B, D, E, and F data are mean ± SEM. Statistical significance was determined using unpaired two-tailed Student’s t-test. ** p < 0.01, *** p < 0.001, **** p < 0.0001

**Suppl 2.**
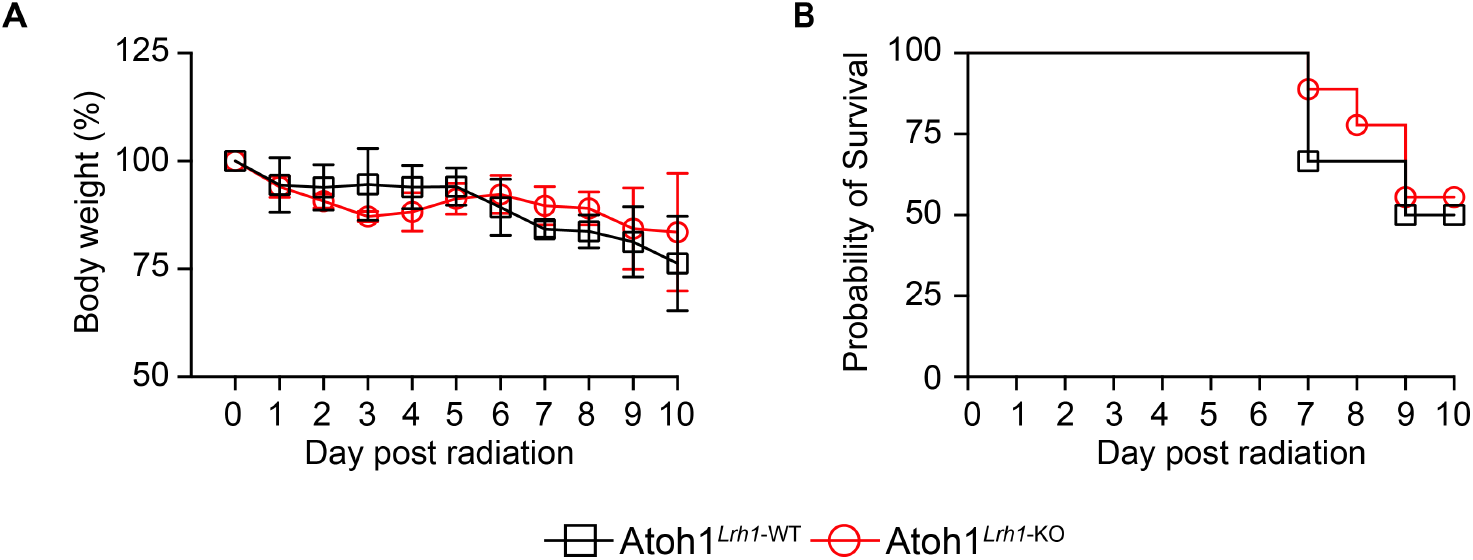
*Atoh1*^*Lrh-1-WT*^ and *Atoh1*^*Lrh1-KO*^ have similar responses to whole body radiation. **(A)** Analysis of mouse body-weight changes after receiving 8Gy-TBI of Lrh1^Atoh1-WT^ and Lrh1^Atoh1-KO^ animals by two-way ANOVA showed no statistical difference between groups. **(B)** Survival rate after 8Gy-TBI between *Lrh1*^Atoh1-WT^ and *Lrh1*^Atoh1-KO^ groups is similar by Kaplan-Meier survival analysis.

**Suppl 3.**
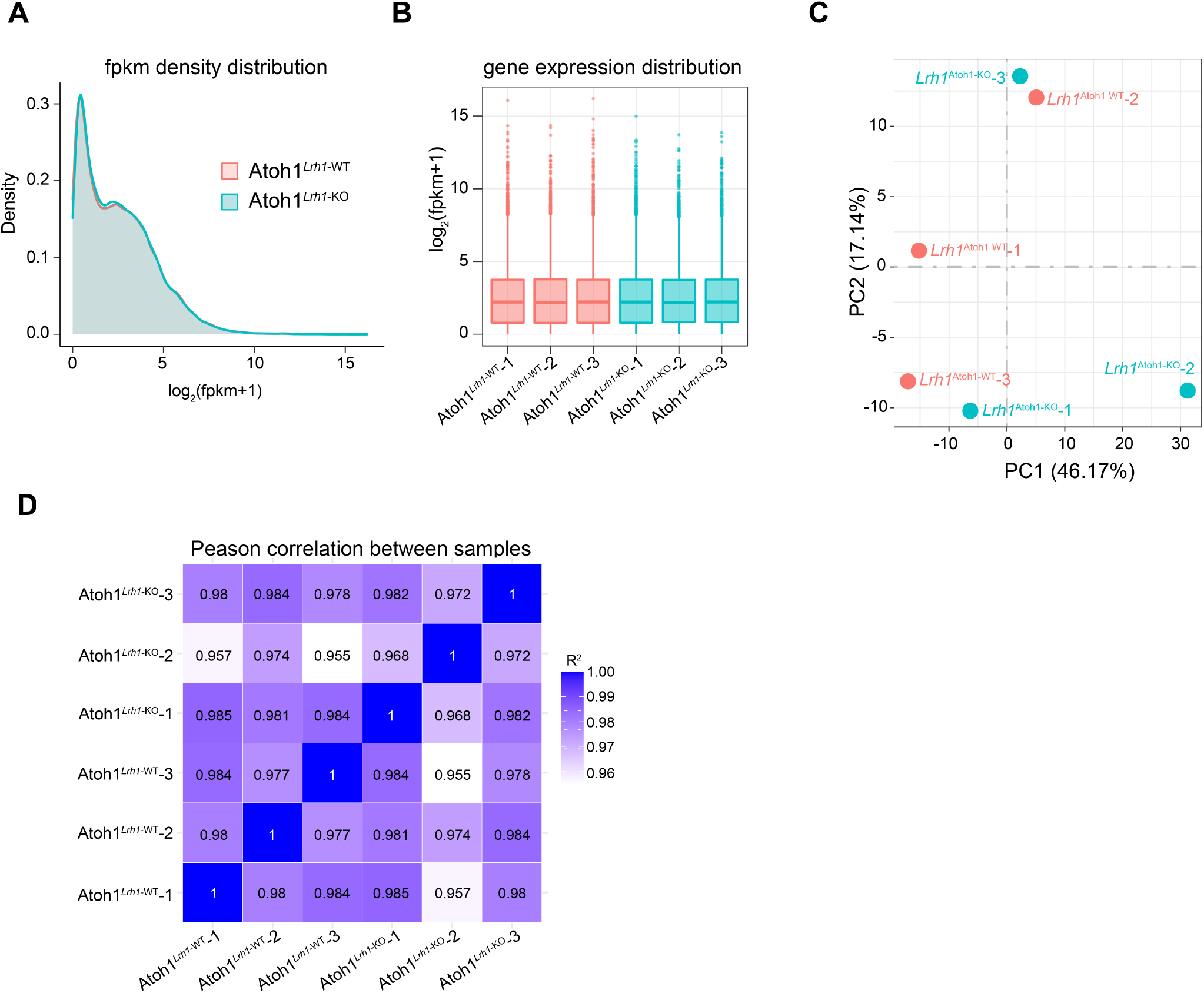
Transcriptomic signature in *Lrh-1* Atoh1-KO overlaps with WT under homeostatic conditions. **(A)** FPKM density distribution plot of *Lrh1*^Atoh1-WT^ and *Lrh1*^Atoh1-KO^ groups. **(B)** Boxplot of gene expression distribution among biological repeats of *Lrh1*^Atoh1-WT^ and *Lrh1*^Atoh1-KO^ animals. **(C)** Principal component analysis revealed limited separation between *Lrh1*^Atoh1-WT^ and *Lrh1*^Atoh1-KO^ groups. **(D)** Peason’s correlation coefficient showed the correlation between different samples of *Lrh1*^Atoh1-WT^ and *Lrh1*^Atoh1-KO^ groups.

**Suppl 4.**
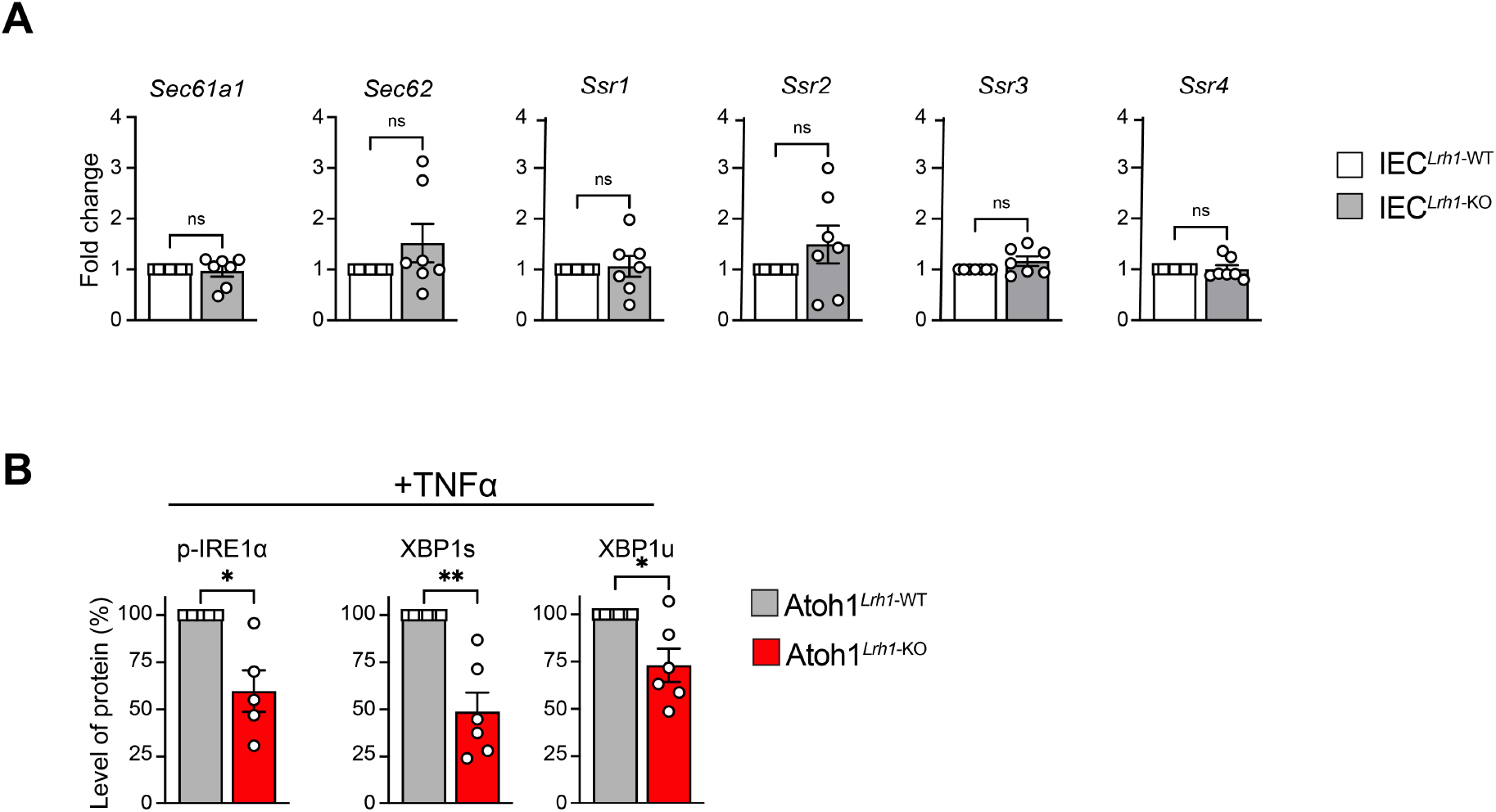
*Lrh1* knockout in Atoh1^+^ lineage led to reduced expression of *IRE1α-XBP1* unfolded protein response genes. **(A)** RT-qPCR results of unsorted organoids showed systematic loss of *Lrh1* did not reduce the gene expression of components involved in peptide transportation across the ER. n=7 per condition. **(B)** Relative quantification of western blot analysis of unsorted *Lrh1*^Atoh1-WT^ and *Lrh1*^Atoh1-KO^ organoids after TNFα treatment in Fig 4E. For both panels, data are presented as mean ± SEM. Statistical significance was determined using unpaired two-tailed Student’s t-test. * p < 0.05, ** p < 0.01

**Suppl 5.**
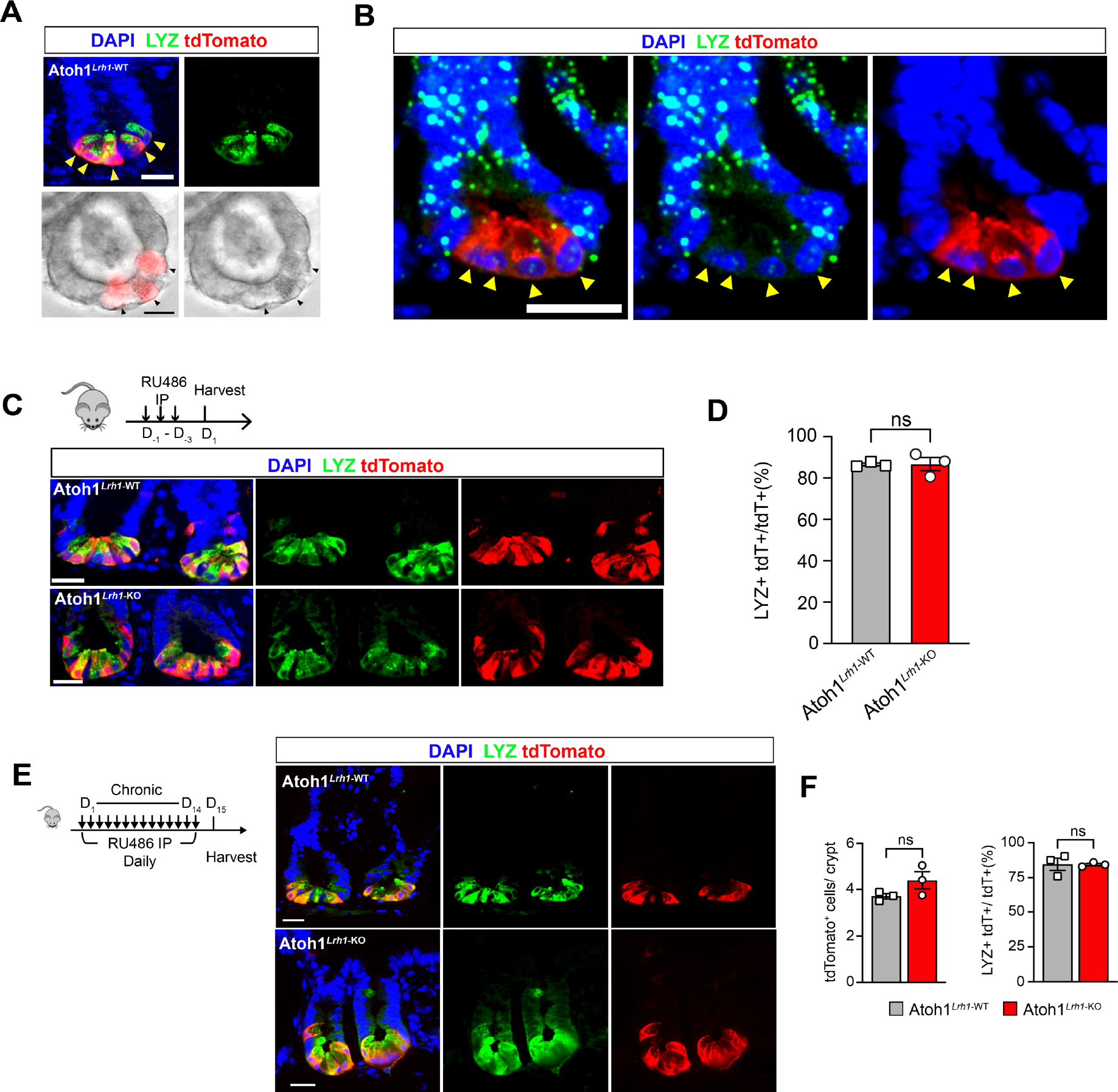
*Lrh1* knockout in Atoh1^+^ lineage impairs the features of Paneth cells. **(A)** Representative images of tdTomato-labeled cells showed the characteristic granule structure of Paneth cells in mouse intestine (top) and intestinal organoid (bottom). **(B)** Representative image showed lower *Lrh1* expression in LYZ^+^ Paneth cells (yellow arrowhead) as compared with TA zone. Scale bar is 20 µm. **(C)** Schematic illustration of the experimental timeline for chronic *Lrh1* deletion in the Atoh1+ lineage and representative images showed Atoh1^CrePGR^-mediated chronic deletion of *Lrh1* caused limited impact on the gross crypt structure. Scale bar is 20µm. **(D)** Quantification of *Atoh1*^CrePGR^-mediated chronic deletion of *Lrh1* neither significantly reduced the number of tdTomato-labeled cells per crypt nor changed the proportion of LYZ+ tdT+ cells per crypt of *Lrh1*^Atoh1-KO^ mice compared to controls. N =3 per condition. **(E)** Schematic illustration of the experimental timeline for chronic *Lrh1* deletion in the Atoh1+ lineage and representative images showed Atoh1^CrePGR^-mediated chronic deletion of *Lrh1* caused limited impact on the gross crypt structure. Scale bar is 20µm. **(F)** Quantification of *Atoh1*^CrePGR^-mediated chronic deletion of *Lrh1* neither significantly reduced the number of tdTomato-labeled cells per crypt nor changed the proportion of LYZ+ tdT+ cells per crypt of *Lrh1*^Atoh1-KO^ mice compared to controls. N =3 per condition. For panels D and F, data are presented as mean ± SEM. Statistical significance was determined using unpaired two-tailed Student’s t-test.

